# Capture-based DNA methylation sequencing facilitates diagnosis and reveals potential pathogenic mechanisms in teratogenic diabetes exposure

**DOI:** 10.1101/172262

**Authors:** Katharina V. Schulze, Amit Bhatt, Mahshid S. Azamian, Nathan C. Sundgren, Gladys Zapata, Patricia Hernandez, Karin Fox, Jeffrey R. Kaiser, John W. Belmont, Neil A. Hanchard

**Affiliations:** Department of Molecular and Human Genetics, Baylor College of Medicine, Houston, TX, USA; Department of Pediatrics, Baylor College of Medicine, Houston, TX, USA; USDA/ARS/Children’s Nutrition Research Center, Baylor College of Medicine, Houston, TX, USA; Department of Obstetrics and Gynecology, Baylor College of Medicine, Houston, TX, USA; Illumina, Inc., San Diego, CA, USA

**Author notes:** **Corresponding Author**: <u>Address:</u> Department of Human and Molecular Genetics, Baylor College of Medicine, One Baylor Plaza, Houston, TX, USA 77030 <u>Email:</u> <u>Tel:</u>.

## Abstract

Diabetic embryopathy (DE) describes a spectrum of birth defects associated with a teratogenic exposure to maternal diabetes *in utero.* These defects strongly overlap the phenotypes of known genetic syndromes; however, the pathogenic mechanisms underlying DE remain uncertain and there are no definitive tests that distinguish the diagnosis. Here, we explore the potential of DNA methylation as both a diagnostic biomarker and a means of informing disease pathogenesis in DE. Capture-based bisulfite sequencing was used to compare patterns of DNA methylation at 2,800,516 sites genome-wide in seven DE neonates and 11 healthy neonates, including five with *in utero* diabetes exposure. DE infants had significantly lower global DNA methylation (ANOVA, Tukey HSD *p*=0.045) than diabetes-unexposed, healthy controls (UH), with multiple sites showing large (mean methylation difference = 16.6%) and significant (*p*<0.001) differential methylation between the two groups. We found that a subset of 237 highly differentially methylated loci could accurately distinguish DE infants from both UH and diabetes-exposed healthy infants (sensitivity 80% -100%). Differentially methylated sites were enriched in intergenic (*p*<3.52×10^-15^) and intronic (*p*<0.001) regions found proximal to genes either associated with Mendelian syndromes that overlap the DE phenotype (e.g. *TRIO*, *ANKRD11*), or known to influence early organ development (e.g. *BRAX1*, *RASA3*). Further, by integrating information on *cis*-sequence variation, we found that 39.3% of loci with evidence for allele-specific methylation also showed differential methylation between DE and controls. Our study suggests a role for aberrant DNA methylation and *cis*-sequence variation in the pathogenesis of DE, and highlights the diagnostic potential of DNA methylation for teratogenic birth defects.

## BACKGROUND

Teratogenesis is the disruption of normal fetal development as a consequence of an environmental exposure. This disruption frequently leads to congenital birth defects in the growing offspring^1^. An estimated 10% of all birth defects are likely attributable to such adverse environmental encounters^2^. Prenatal exposure with a teratogenic agent can not only be a consequence of maternal medications or substance abuse^3-5^, but also of maternal health. Pregestational diabetes is a well-known cause of congenital malformations and infants of diabetic mothers (IDMs) are 2- to 4-fold more likely to develop birth defects compared to the general population^6-10^. Likewise, maternal obesity and gestational diabetes (GDM) are associated with increased risks of birth defects^11-14^. As a consequence, approximately 4-9% of children prenatally exposed to maternal diabetes are born with diabetic embryopathy (DE)^6-10^ – a phenotypic spectrum of birth defects that classically includes sacral agenesis and related spine defects, complex congenital heart defects, central nervous system (CNS) malformations, and limb abnormalities. These defects often co-occur as so-called ‘multiple congenital anomalies’ and have substantial phenotypic overlap with recognizable genetic and genomic syndromes^14^. In clinical practice, this poses a difficult diagnostic problem, as the presence of multiple birth defects triggers a broad and often expensive diagnostic odyssey aimed at providing parents with information regarding recurrence risk, future medical management, and recommendations for disease screening. However, as there is presently no definitive diagnostic test for DE, diagnostic evaluations in such contexts are usually futile. The diagnosis of “infant of a diabetic mother”, therefore, relies almost entirely on observing congenital malformations in an infant with a history of maternal diabetes and excluding plausible alternative, usually genetic, diagnoses.

The teratogenic mechanisms underlying maternal diabetes remain elusive, but the likelihood of congenital anomalies following *in utero* maternal diabetes exposure appears to correlate positively with first trimester maternal blood glucose levels^15,16^. DNA methylation, most commonly found at cytosines in cytosine-guanine dinucleotides (CpG), is a malleable regulatory epigenetic feature that can reflect *in utero* exposures to environmental stimuli, such as maternal nutrition^17^. It has thus been hypothesized that maternal diabetes in humans could disrupt normal epigenetic patterns, resulting in aberrant gene function during embryogenesis, and ultimately leading to complex malformations in the fetus. For instance, in a recently published murine model, oxidative stress induced by maternal hyperglycemia led to increased activity of the DNA methyltransferase Dnmt3b, which in turn resulted in DNA hypermethylation and inhibited embryonic expression of *Pax3* - a gene crucial for neural tube and heart development^18^. Most studies of DNA methylation in IDMs, however, have focused on epigenetic changes that could influence the risk of later-life obesity and diabetes in these offspring. These studies have assessed the effects of GDM on DNA methylation in placental tissue and umbilical cord blood in exposed, yet phenotypically healthy progeny and have reported disrupted methylation patterns in both candidate gene analyses^19-22^ and genome-wide array-based scans^23-26^. Using array and methylated DNA immunoprecipitation (MeDIP-chip) technologies, epigenome-wide studies on peripheral blood of older offspring without congenital malformations (ranging from young adolescents to adults) also showed altered DNA methylation patterns in individuals with *in utero* exposure to gestational^27^, type 1 (T1DM)^28^, and type 2 diabetes mellitus (T2DM)^29^. These studies, however, provided only a restricted view of methylation following diabetic pregnancies and none of them assessed methylation in the context of DE.

Given this gap in knowledge, we used genome-wide targeted enrichment bisulfite sequencing to assess DNA methylation across the genome in neonates with congenital malformations and a history of *in utero* diabetes exposure. Buccal epithelial cell samples were obtained for DNA isolation in order to avoid confounding by cell composition differences in peripheral blood^30^. Our starting premise was that teratogenic effects incurred during early embryogenesis will be substantial and affect all germ layers. Given the fidelity of mitotic inheritance of DNA methylation, we hypothesized that traces of these effects would still be evident in differentiated tissues at and around the time of birth. The resulting patterns of DNA methylation were compared with those of healthy infants, who had not been exposed to maternal diabetes (unexposed, healthy; UH), and maternal diabetes-exposed newborns without overt evidence of a congenital clinical phenotype (exposed, healthy; EH). In so doing, we aimed to evaluate the diagnostic potential of the methylation and possibly gain insight into the underlying pathogenesis.

## MATERIALS AND METHODS

### Participants

Participants were recruited through Texas Children’s Hospital in Houston, Texas, as part of a larger study of the effects of maternal diabetes and anti-epileptic drug consumption on offspring DNA methylation. The study was reviewed and approved by the Baylor College of Medicine Institutional Review Board. In total, 22 neonates were enrolled (**Table 1**) – 15 were born to diabetic mothers, of whom nine were diagnosed with DE. The remaining six diabetes-exposed infants did not have any clinical stigmata of diabetes exposure, including respiratory distress or metabolic abnormalities such as hypoglycemia. All but three of our cohort were carried to term (two UH, one EH), and all births occurred after 36 completed weeks of gestation (median: 38.3 weeks, range 36.4 to 40.1 weeks; **Table 1**). Our study design was inclusive of all types of maternal diabetes, however, GDM and T2DM were the most commonly observed (**Table 1**). The congenital malformations found in these infants, the genetic tests undertaken to eliminate known genetic disorders as alternative diagnoses, as well as infant and mother demographics are listed in **Supplementary Table 1**. At the time of enrollment, mothers of infants with congenital malformations had significantly higher body mass indices (BMIs) compared to healthy mothers (*p*=0.017, Tukey HSD; **Table 1**), making maternal obesity an unavoidable confounder of diabetes exposure. Maternal blood glucose concentrations, assessed by glycated hemoglobin (HbA1c) levels, were only available for one UH control sample, thus impeding any statistical comparisons; nonetheless, the two highest HbA1c measurements were observed in mothers of infants with DE (**Supplementary Figure 1**). This trend is consistent with previously reported positive correlations between maternal blood glucose levels and the risk of associated congenital malformations^16,15^. The American Congress of Obstetrics and Gynecology recommends screening for diabetes in pregnancy, and chart review of all UH mothers showed that none had GDM. We did not find any significant differences in maternal age or infant birth weight (**Table 1**).

**Table 1:**
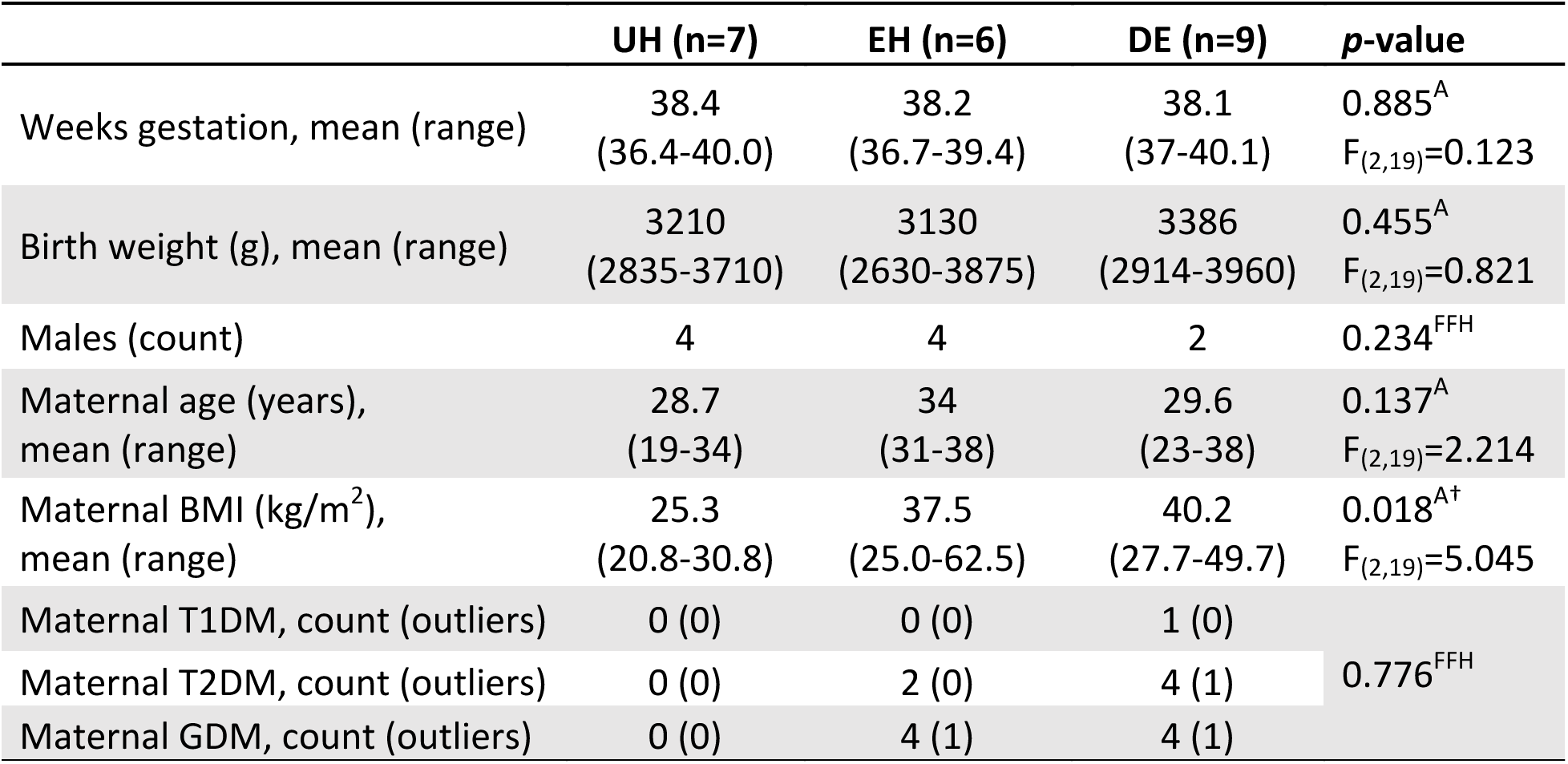
Maternal and infant characteristics. Body mass index (BMI) at enrollment; UH=diabetes unexposed, healthy infants; EH=diabetes exposed, healthy infants; DE=infants with diabetic embryopathy; T1DM=type 1 diabetes mellitus; T2DM=type 2 diabetes mellitus; GDM=gestational diabetes mellitus; ^A^=one-way ANOVA; ^FFH^=Fisher-Freeman-Halton exact test; †Tukey’s HSD adjusted *p*-value=0.017.

|  | UH (n=7) | EH (n=6) | DE (n=9) | p-value |
| --- | --- | --- | --- | --- |
| Weeks gestation, mean (range) | 38.4<br>(36.4-40.0) | 38.2<br>(36.7-39.4) | 38.1<br>(37-40.1) | 0.885<sup>A</sup><br>F<sub>(2,19)</sub>=0.123 |
| Birth weight (g), mean (range) | 3210<br>(2835-3710) | 3130<br>(2630-3875) | 3386<br>(2914-3960) | 0.455<sup>A</sup><br>F<sub>(2,19)</sub>=0.821 |
| Males (count) | 4 | 4 | 2 | 0.234<sup>FFH</sup> |
| Maternal age (years),<br>mean (range) | 28.7<br>(19-34) | 34<br>(31-38) | 29.6<br>(23-38) | 0.137<sup>A</sup><br>F<sub>(2,19)</sub>=2.214 |
| Maternal BMI (kg/m<sup>2</sup>),<br>mean (range) | 25.3<br>(20.8-30.8) | 37.5<br>(25.0-62.5) | 40.2<br>(27.7-49.7) | 0.018<sup>A†</sup><br>F<sub>(2,19)</sub>=5.045 |
| Maternal T1DM, count (outliers) | 0 (0) | 0 (0) | 1 (0) | 0.776<sup>FFH</sup> |
| Maternal T2DM, count (outliers) | 0 (0) | 2 (0) | 4 (1) |  |
| Maternal GDM, count (outliers) | 0 (0) | 4 (1) | 4 (1) |  |

### Sample collection and sequencing

Buccal epithelial cells were collected within ten days of birth using the Oragene Discover (OGR-250) DNA collection kit (DNA Genotek Inc., Ottawa, Ontario, Canada), modified to obtain a cumulative sample of the inner buccal cheek by doing multiple (10) passes with multiple (5) swabs. DNA was extracted according to the manufacturer’s instructions. In order to quantify DNA methylation epigenome-wide, the SeqCap Epi CpGiant Enrichment Kit (Roche NimbleGen, Inc., Madison, Wisconsin, USA) was used. This is a targeted methylation capture panel designed to assess ~5.5 million methylation sites across the genome. Library preparation, bisulfite conversion, targeted enrichment, and amplification were performed as outlined in the SeqCap Epi CpGiant Enrichment Kit protocol. Adapters from the SeqCap Adapter Kit A were used in combination with SeqCap EZ HE-Oligo Kit A barcode blockers (both from Roche NimbleGen, Inc., Madison, Wisconsin, USA) to index samples. The captured DNA fragments were amplified, then sequenced on the Illumina HiSeq 2000 platform using 100 bp paired-end reads, and multiplexed to have three samples per lane. Samples were sequenced in two batches: the first (discovery) batch comprised three UH, one EH, and four DE infants, and the second (replication) batch consisted of three UH, four EH, and three DE samples, excluding outliers (see below).

### Post-sequencing processing and quality control

De-multiplexed BAM files were processed using Bismark (version 0.12.3)^31^, including (1) alignment of sequence reads to the hg19 reference genome, (2) elimination of PCR duplicates, (3) removal of five bases from 5’ read ends in order to reduce known technical biases, and (4) quantification of the methylation level at each targeted CpG locus. The average mapping efficiency was 77.15%, with a mean of 45,140,004 uniquely aligned reads per sample (median: 46,341,251, range: 34,344,965 to 52,937,052). CpG loci overlapping common SNPs (MAF ≥1% in hg19 dbSNP build 144) were discarded from subsequent analyses. The average capture efficiency after deduplication - defined here as the number of reads overlapping targeted CpG loci divided by the total number of sequencing reads - was 65%. The median coverage at all CpG sites targeted by the enrichment kit was 20X (range: 1X to 13,698X). Downstream analyses were performed using only those CpG sites with ≥10X coverage in every sample in a given analysis (median: 26X). Potential batch effects were evaluated using one-way ANOVA on the medians of the distributions as well as quantro software (1,000 data permutations; version 1.4.0)^32^. There was no strong evidence of batch effects, nor were there other biases noted in the distribution of cases, infant demographics or maternal demographics (**Supplementary Table 2**).

To confirm that our DNA samples were in fact derived from buccal epithelial cells, we isolated available CpG sites from our data set that are needed for the online Horvath DNA methylation age calculator (accessed Nov. 22, 2016)^33^, which provides probabilities for likely tissues of origin based on tissue-specific methylation patterns. Percent methylation values were scaled to proportions in order to emulate beta values - the measure of methylation calculated in array analyses and required by the tool. Based on the tissue prediction algorithm, all samples were confirmed to be primarily buccal (epithelial) in origin (**Supplementary Table 3**).

Given the breadth of clinical phenotypes evident in the cohort, a principal component analysis (PCA) and hierarchical clustering of samples using all sites with ≥10X coverage across all individuals was performed. Four outlier samples – one UH, one EH, and two DE infants (**Supplementary Figures 1A and B**) – were removed following this analysis. This is conservative in that it assumes that most CpG sites are not differentially methylated comparing cases and controls. After outlier removal, a total of 3,108,549 CpG sites remained in DE infants, 3,176,002 in UH infants, and 3,791,204 in EH infants; 2,760,543 were shared among all groups; 2,800,516 sites were shared between UH and DE samples.

### Differential methylation analysis

Single site differential methylation analysis using a logistic regression model without covariates, was performed using methylKit (version 0.9.5)^34^, after normalization for coverage using the same tool. In an effort to reduce the number of false positive results, all sites with nominal p-values <0.001 were permuted 1,000 times with replacement by randomly switching case/control labels. In order to give greater confidence to our findings, we took advantage of the known correlation between methylation of adjacent CpG dinucleotides to cluster sites into bins of multiple high-confidence differentially methylated loci – focusing on sites at which ≤10 permutations produced an equally small or smaller p-value than the original analysis. All high-confidence sites within a bin were required to be separated from neighboring sites by no more than 1 kb and share the same direction of effect. Bins that did not contain at least one high-confidence differentially methylated site with an absolute difference (Δ) in percent methylation ≥10 (i.e. ≥10 absolute percentage points – pp) were excluded, as were those that did not contain at least one differentially methylated CpG site (p<0.05, ≤10/1,000 better or equal permutations) using methylSig (version 0.4.1) ^35^ – an alternative method of assessing differential methylation based on a beta-binomial approach.

### Joint batch differential methylation analysis

We initially performed differential methylation analysis as outlined above using all DE and UH samples, focusing on the largest bins (containing ≥3 high-confidence sites). On average, in this analysis, coverage normalization resulted in 0.53 pp difference (i.e. <1% absolute difference in magnitude) in methylation (range: 0 to 3.79 pp) at high-confidence differentially methylated CpG loci from candidate bins.

To test classification using methylation in our joint analysis, we used four-fold cross-validation with the default settings of WEKA’s “Logistic” classifier (version 3.6.13)^36^ and ten repetitions on the joint batch data set. Four-fold cross-validation involves first dividing the data set into four bins of equal size; next one bin is withheld to serve as test data set and the classifier is trained on the remaining bins. Accuracy is determined by the degree of concordance between the reported and classified sample labels. The process is then repeated ten times, each time reallocating the samples into new bins.

### Classification

In order to assess the utility of methylation patterns for correctly assigning cases and controls when blinded to sample labels, we repeated the differential methylation analysis using only the discovery batch, with two exceptions, both designed to account for the smaller sample size. First, we imposed a more stringent bin size of ≥7 high-confidence sites (**Supplementary Figure 3**); second, in the methylSig analysis, we decreased alpha to 0.01 for the retention of bins containing at least one CpG locus which passed that threshold of differential methylation. Classification was performed using WEKA’s logistic regression model on the high-confidence sites found in the largest bins shared between the discovery and the test batches. As a secondary approach, we also applied a classification model previously used to distinguish individuals based on their DNA methylation patterns^37^. This entailed first calculating the median DNA methylation values at the selected loci for the case and control samples in the discovery set, and then determining Pearson correlation coefficients for the corresponding CpG sites of each sample in the test set. A sample would then be classified as case or control depending upon which coefficient (case or control) was larger.

Coverage normalization during data processing altered percent methylation at classifier loci by an average of 0.57 pp (i.e. <1% absolute difference in magnitude; range: 0 to 4.55 pp) in the DE and UH comparison, and by 0.34 pp (range: 0 to 3.33 pp) in the DE and EH comparison. These observations suggested that our results were unlikely to be the consequence of coverage normalization artifacts.

### Gene ontology and pathway enrichment analysis

RefSeq genes overlapping or within 1 kb of our candidate regions were taken as candidate genes. ConsensusPathDB’s over-representation analysis (accessed Feb. 2, 2017)^38^ was employed to identify enriched pathways and functional annotations from these candidates. RefSeq genes overlapping or within 1 kb of the CpG loci targeted by the enrichment kit were used as background. All pathway database options except ‘SMPDB’ and ‘PharmGKB’ were included and default parameters were used for each analysis. Gene ontology analysis included level 2-5 categories with default parameters.

### Allele-specific methylation analysis

BisulfiteGenotyper in the Bis-SNP package (version 0.82.2)^39^, which is built upon the GATK framework, was used to call single nucleotide variants (SNVs) within 100 bp (the length of a sequencing read) of all targeted CpG sites. The algorithm was used with the hg19 reference genome and dbSNP build 144. The resulting variant files, which included all confident sites, were merged into a single file using GATK’s CombineVariants function (version 2.3-9-ge5ebf34)^40^. Filtering steps included the removal of loci with overlap of variant and targeted CpG sites, SNVs at bases affected by bisulfite treatment (cytosines and guanines), and minus strand cytosines. The latter were omitted to avoid the overrepresentation of each CpG locus, as plus and minus strand cytosines of the same CpG dinucleotide are closely correlated. The comparison of percent methylation at heterozygous SNVs in a single individual, was limited by the small number of available high coverage reads spanning both SNVs and CpGs and was not included in our analysis. Instead, allele-specific methylation (ASM) was assessed at SNV-CpG pairs with at least 10X coverage that were observed in at least two homozygous individuals for each reference and alternate allele. Percent methylation and coverage were averaged across each allelic state. Coverage discrepancies were limited and not statistically significant at SNV-CpG pairs of interest (*p*>0.05, two-tailed t-test; **Supplementary Figure 4**).

### Statistical analysis

All statistical analyses were performed in R. Quantitative demographics as well as batch percent methylation were compared using one-way ANOVA; statistically significant ANOVA comparisons between three groups were followed by a Tukey HSD test to determine significant pair-wise differences. Categorical demographics were assessed between two or three groups by two-sided Fisher’s exact and Fisher-Freeman-Halton tests, respectively. Enrichment and depletion of gene annotations in data subsets were evaluated against the annotations of all tested sites using a hypergeometric test. The threshold of significance was set at *p*<0.05 for the above analyses.

### Data availability

The datasets generated during and/or analyzed during the current study are available from the corresponding author on reasonable request.

## RESULTS

### Depletion of DNA methylation characterizes infants with diabetic embryopathy

Of the 22 infants enrolled, nine were given a diagnosis of DE based on the presence of maternal diabetes during pregnancy and the observation of congenital structural defects consistent with that diagnosis; six IDMs were completely healthy at birth (EH), and seven were healthy infants born to healthy mothers (UH). After removing four outliers (2 DE, 1 EH, and 1 UH; **see methods**), we evaluated buccal DNA methylation at a total of 2,760,543 CpG sites shared between all infant groups. Global mean percent DNA methylation levels were significantly lower in DE infants compared to UH controls (42.6% versus 43.2%; *p*=0.045, Tukey HSD; **Figure 1A**). DNA methylation throughout the genome typically follows a bimodal distribution in which most CpG loci are either methylated or unmethylated, with fewer sites having intermediate methylation^41^. In order to assess differences between these three methylation categories, CpG sites were separated into bins of low (<30%), intermediate (30-70%), and high (>70%) DNA methylation using UH control samples as a reference. There was no statistically significant difference in global methylation between infant groups in the low-(*p*=0.298, F_(2, 15)_=1.313, one-way ANOVA; **Figure 1B**) or intermediate-(*p*=0.142, F_(2, 15)_ =2.228, one-way ANOVA; **Figure 1C**) methylation bins; however, among DE infants, we observed significant DNA hypomethylation at highly methylated sites when compared to UH controls (*p*=0.011, Tukey HSD; **Figure 1D**). These results suggested that the global reduction of DNA methylation seen in DE neonates is largely a consequence of reduced methylation at CpG sites that are highly methylated in healthy, un-exposed infants.

**Figure 1:**
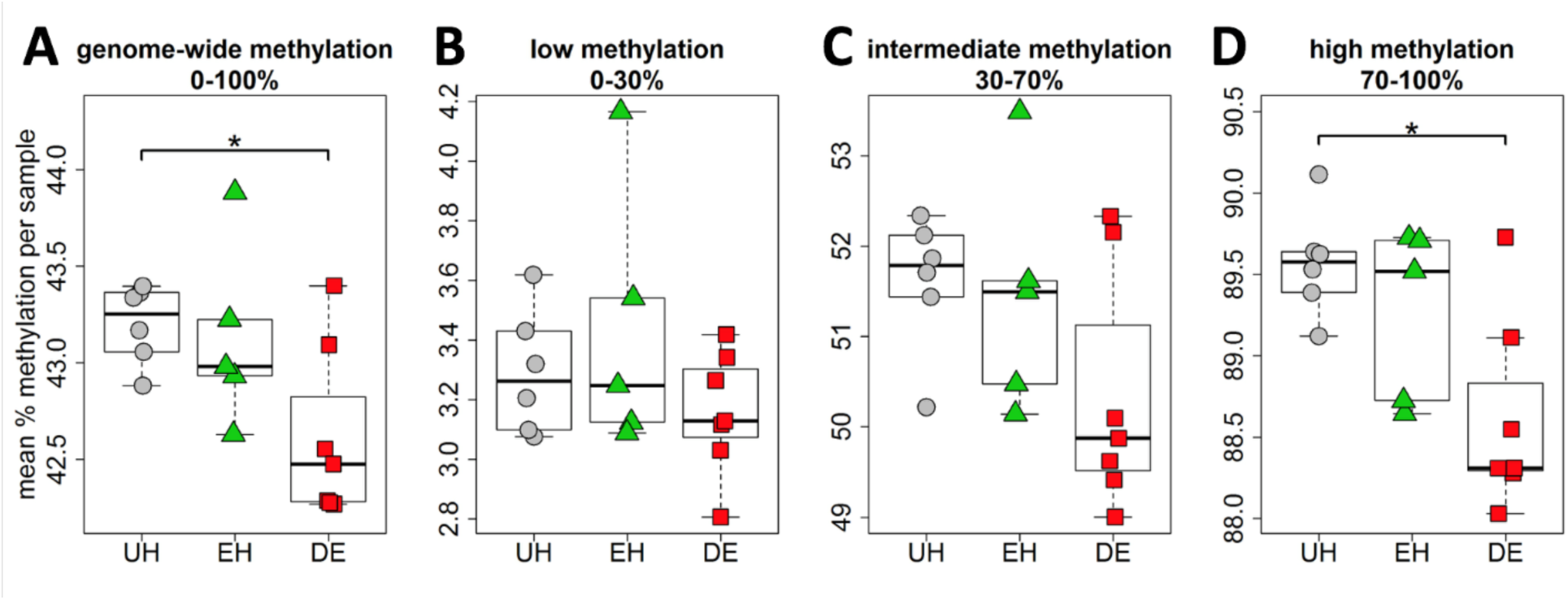
Infants with diabetic embryopathy (DE) show a global loss in DNA methylation compared to unexposed, healthy controls (UH). All boxplots are comparisons of mean sample percent methylation across the number of sites in each category. (**1A**) 2,760,543 sites shared among DE, UH, and diabetes exposed, yet healthy infants (EH); (**1B**) 1,362,136 (49.3%) sites with mean methylation <30% in UH infants; (**1C**) 276,215 (10.0%) sites with mean methylation 30-70% in UH infants; (**1D**) 1,122,192 (40.7%) sites with mean methylation >70% in UH infants. ^∗^Tukey HSD *p*<0.05.

### Differentially methylated sites can distinguish clinical outcomes among diabetes-exposed and unexposed infants

In light of the epigenetic changes identified in DE infants at the global level, we next considered the utility of DNA methylation as a biomarker of the maternal diabetes exposure. A linear regression model was used to evaluate methylation differences between UH and DE. In total, 13,239 (0.5%) high-confidence sites showed evidence of differential methylation (*p*<0.001) and surpassed our stringent, permutation-based cut-off (**see methods**). We then leveraged the anticipated correlations between neighboring CpG loci to further eliminate spurious associations by clustering high-confidence sites into non-overlapping bins. Differentially methylated sites fell into 237 high-confidence bins (≥3 high-confidence differentially methylated sites per candidate bin, each containing a CpG site with *p*<0.05 using our secondary method; **see methods**). Consistent with our global analysis, the majority (87.8%) of these bins showed hypomethylation across clustered sites (**Supplementary Table 4**). The mean absolute difference in percent methylation across all 1,010 high-confidence CpG sites in these bins was 16.6% (SE 0.3%). Using PCA on these high-confidence sites, we found that UH samples clustered separately and distinctly from infants with DE (**Figure 2A**). Moreover, when we repeated the same analysis, this time including EH infants, we observed a distinct clustering of EH infants on the first principal component that was separate from, and intermediate to, DE cases and UH controls (**Figure 2B**). The resulting heat map of differentially methylated sites further bolstered this observation – EH individuals demonstrated a mixed, intermediate, pattern to either UH or DE, with most sites having a comparable magnitude of differential methylation (**Figure 2C**). Four-fold cross-validation (**see methods**) correctly categorized samples with 100% accuracy, suggesting that the differential methylation patterns are distinct enough to be exploited for classification.

**Figure 2:**
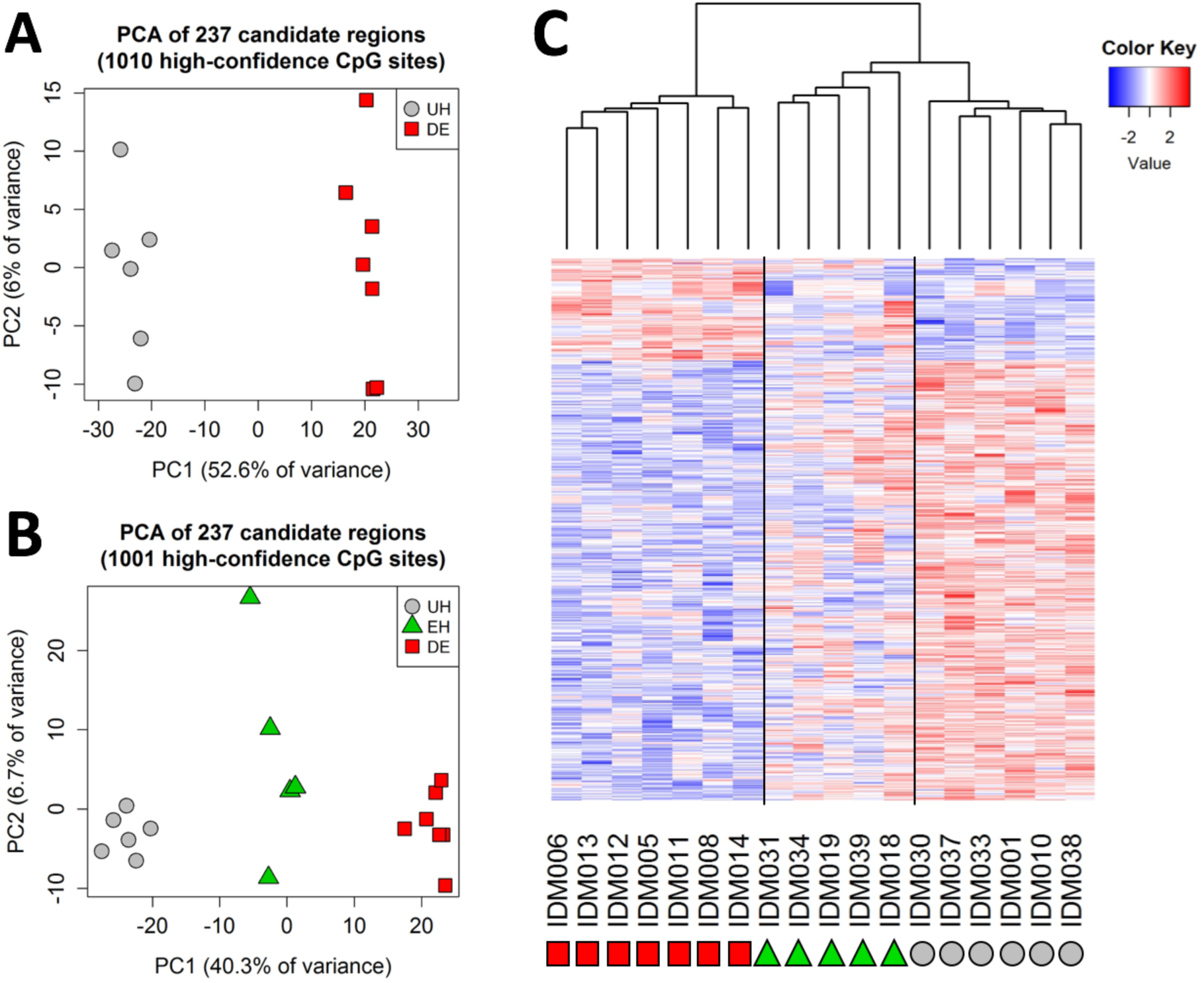
Distinct DNA methylation profile in DE cases compared to controls. Each analysis was based on all sites found in candidate bins with ≥10X coverage and non-zero variance across samples. (**2A**) Principal component analysis (PCA) based on 1,010 high-confidence differentially methylated CpG sites shared across all DE and UH infants. (**2B**) PCA based on 1,001 high-confidence differentially methylated CpG sites shared across all DE, UH, and EH infants. Difference in numbers between 2B and 2A is the result of coverage adequacy in EH samples. (**2C**) Heat map of scaled and centered percent methylation values corresponding to 2B, with each row representing one high confidence differentially methylated site, and each column a sampled individual.

We then further explored the diagnostic potential of these methylation patterns by assessing whether highly differentially methylated sites could be used to blindly classify clinical status. We did not find evidence for significant batch effects between our first (discovery) and second (replication) batches (*p*=0.544, F_(1, 11)_=0.392, one-way ANOVA; quantro permutation *p*=0.597; **see methods**; **Supplementary Table 2**). Therefore, we started by evaluating differential methylation in our discovery batch of 4 DE and 3 UH samples. Of the 3,092,753 shared CpG sites analyzed, 13,731 high-confidence sites (0.4%) showed evidence of differential methylation (*p*<0.001) and surpassed our stringent permutation-based cut-off. These sites were further filtered to 15 high-confidence bins, containing at least one site with ≥10% absolute difference in methylation, using the same criteria as the joint analysis, with the exception of necessitating seven, instead of three, differentially methylated loci (**see methods**; **Supplementary Figure 3**). High-confidence bins had an average absolute difference in methylation of 28.6 percentage points (SE 2.0) between DE and UH infants, and again the majority (73.3%) were hypomethylated in DE infants.

Using the 156 CpG sites in our high-confidence bins that had adequate coverage in our test batch, our logistic regression model (**see methods**) was able to classify the remaining six UH and DE test samples with 100% accuracy (6/6). To avoid overfitting, we applied a secondary classifier built on the correlation of methylation values from each test sample with either case or control DNA methylation values from the discovery batch (**see methods**). Using this approach, we were able to distinguish between UH and DE samples in 5/6 (83.3%) instances (**Figure 3A**). To investigate how well EH infants would be grouped using a classifier trained to distinguish DE from UH individuals, we applied both methods using the 155 CpG sites shared by all infants. The logistic regression model classified 2/5 (40%) of EH samples as “DE”, thereby performing no better than random; however, the correlation-based model labeled 100% of EH samples as “DE”, suggesting that this method might be better for the distinction between diabetes exposed and unexposed individuals.

**Figure 3:**
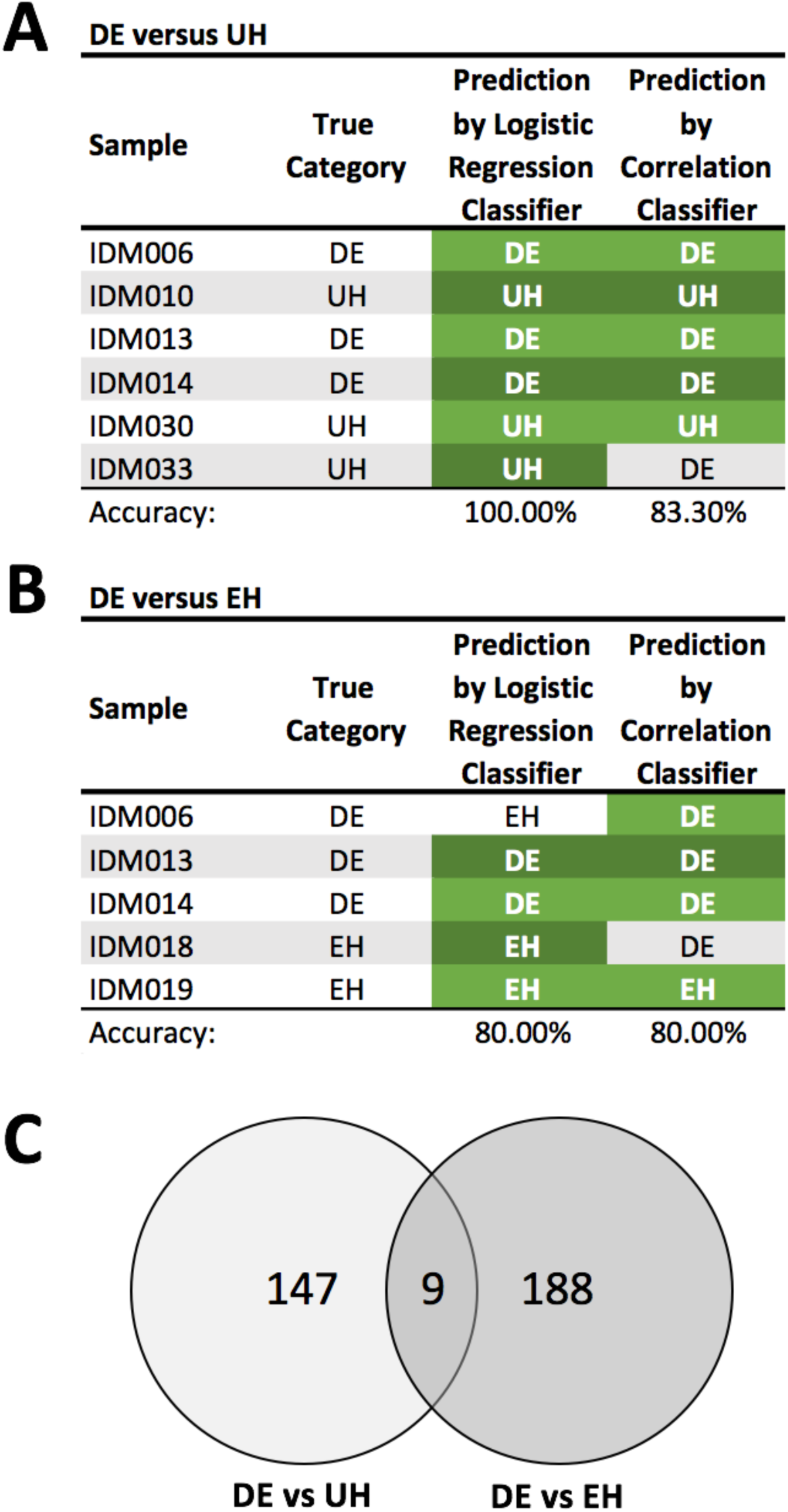
Batch classification results. Predictions in green were classified correctly in the comparisons between DE and UH (**3A**), and EH (**3B**) samples. Of the CpG loci used to distinguish between sample groups in each comparison, nine were present in both (**3C**).

Not all pregnancies exposed to maternal diabetes result in birth defects; therefore, we evaluated the potential to discern malformation from exposure by comparing the two groups with *in utero* exposure to maternal diabetes – DE and EH – in the same classifier analysis. Since most EH samples were sequenced in the second batch, we carefully matched them on the basis of demography such that we had three infants in the discovery cohort and two in the replication set (test of batch effects: *p*=0.769, F_(1, 10)_=0.091, one-way ANOVA; quantro permutation *p*=0.673; **Supplementary Table 2**). Differential methylation analysis and permutation testing revealed 19,139 (0.6% of 3,197,718 loci) high-confidence differentially methylated sites, of which 197 fell into 21 high-confidence bins. In contrast to the previous comparison, however, fewer than half of these bins were hypomethylated in DE relative to EH individuals (10/21, 47.6%).

Both classification methods correctly identified 80% (4/5) of samples, although the misidentified individuals differed between classifier methods (**Figure 3B**). When we looked at the overlap between loci used to distinguish infants with DE from both UH and EH neonates, nine differentially methylated sites were characteristic of the malformation group (**Figure 3C**). Somewhat surprisingly, all sites were contained in the same bin and were hypermethylated in DE infants. This region is less than 4.5 kb upstream of *CSF3* (chr17: 38,167,214 – 38,167,428) – a granulocyte colony stimulating factor that, in rodents, has been shown to protect against left ventricular remodeling and cardiac myocyte apoptosis after myocardial infarction^42^.

### IDM-related differential methylation occurs at non-promoter sites of developmental genes

To further characterize the changes in DNA methylation and to gauge how these might relate to the clinical presentation of diabetic embryopathy, we interrogated the gene content of the differentially methylated sites from the initial joint analysis. Among the binned loci of interest, we found an expansion of the previously mentioned, DE-characteristic region upstream of *CSF3*, which was second in the number of binned CpG sites only to another hypermethylated region encompassing two microRNAs of unknown function, *MIR3648* and *MIR3687* (**Table 2**, **Figure 4A**, and **Supplementary Table 4**). In this list were also several genes associated with genetic syndromes whose phenotypic spectrum closely overlaps the clinical features of DE (**Table 2** and **Supplementary Table 4**); this included *ANKRD11*, the causal gene in KBG syndrome (MIM #148050; **Figure 4B**), which includes spinal and digit malformations as well as occasional heart defects; *B3GNT1*, associated with Walker-Warburg syndrome (MIM #615287); *BRF1*, which results in the characteristic CNS and skeletal abnormalities of cerebellofaciodental syndrome (MIM #616202); *CACNA1C*, which gives rise to the cardiac and digit abnormalities noted in Timothy syndrome (MIM#601005), and *ZBTB20* which causes the large stature noted as part of Primrose syndrome (MIM #259050). We also observed significant differential methylation at other genes that result in DE-like features when knocked down in mouse models, including *BARX1* (embryonic lethal with cleft palate^43^; **Figure 4C**) and *RASA3* (abnormal embryogenesis including abnormal vascular endothelial cell development^44^).

**Table 2:**
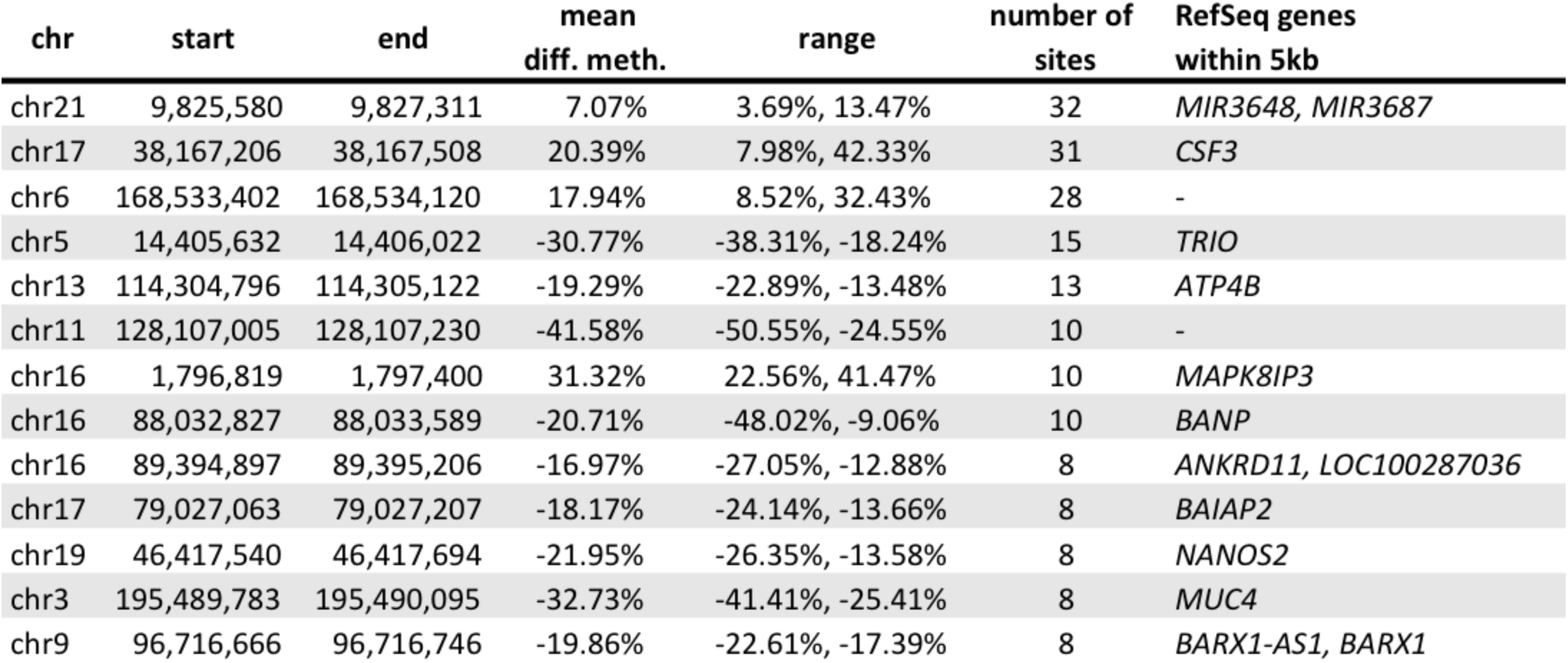
Candidate binned regions with the most high-confidence differentially methylated CpG sites. A complete list of all 237 CpG bins can be found in Supplementary Table 3.

**Figure 4:**
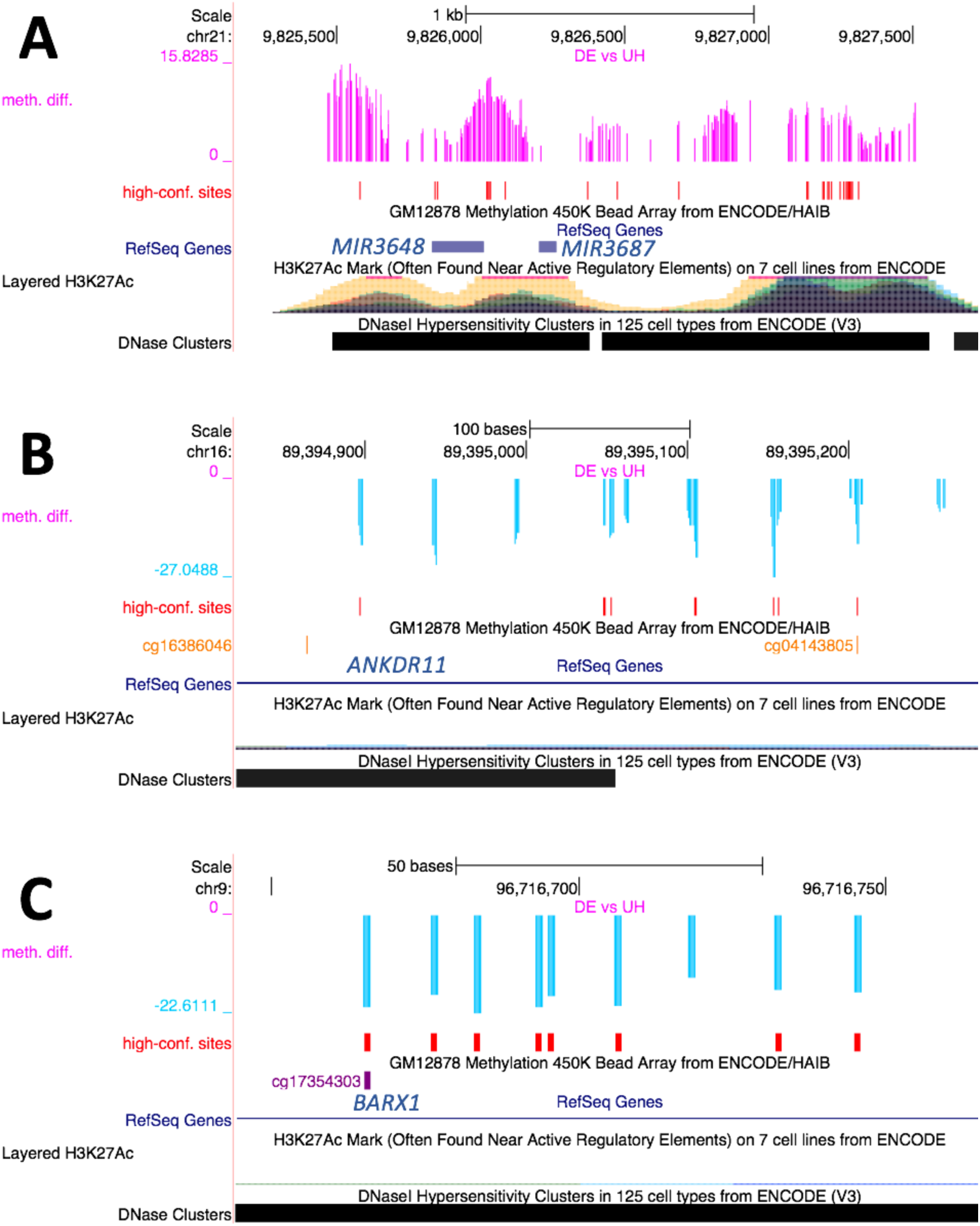
Differentially methylated sites in candidate bins, showing *MIR3648* and *MIR3687* (**4A**), *ANKRD11* (**4B**), and *BARX1* (**4C**). Differences in methylation are found in the “meth.diff” track. Hypomethylation in DE infants is denoted with negative values (blue bars), while positive values (pink bars) indicate hypermethylation. High-confidence differentially methylated sites are marked in red.

In order to get a broader sense of the genes implicated in our analysis, we performed gene ontology and pathway enrichment analyses using all genes within 1 kb of binned regions (n=176). Out of six enriched pathways (nominal *p*≤0.01), two were related to cardiac and neuron function (*Phase 2* - *plateau phase* and *DCC mediated attractive signaling*; **Supplementary Table 5**) - both major systems affected in DE. The former included three of our candidate genes, *CACNA1C*, *CACNG6*, and *KCNQ1*, all of which encode ion-gated channels, while the latter encompasses *PTK2*, a tyrosine kinase, and *TRIO*, a serine/threonine kinase and guanine exchange factor associated with autosomal-dominant mental retardation (MIM #617061). Among the 24 gene ontology terms for which our candidate gene set was enriched, ten related to basic cellular and organismal development including actin function, ion binding, cell development, and nervous system function and development (**Table 3**).

**Table 3:** Enrichment of development-related gene ontologies in candidate genes; ontologies with enrichment *p* ≤0.01 are shown.

| Enriched gene ontologies (GO ID) | p-value | q-value |
| --- | --- | --- |
| actomyosin (GO:0042641) | 0.00001 | 0.00049 |
| receptor signaling complex scaffold activity (GO:0030159) | 0.00081 | 0.03882 |
| contractile actin filament bundle (GO:0097517) | 0.00081 | 0.00701 |
| actin cytoskeleton (GO:0015629) | 0.00082 | 0.00701 |
| actin filament bundle (GO:0032432) | 0.00108 | 0.02530 |
| metal ion binding (GO:0046872) | 0.00180 | 0.04317 |
| neurofilament (GO:0005883) | 0.00224 | 0.03512 |
| pyridine nucleotide metabolic process (GO:0019362) | 0.00268 | 0.22828 |
| cation binding (GO:0043169) | 0.00281 | 0.13482 |
| transcriptional repressor complex (GO:0017053) | 0.00365 | 0.17878 |
| pyridine-containing compound metabolic process (GO:0072524) | 0.00368 | 0.43572 |
| regulation of meiotic cell cycle (GO:0051445) | 0.00408 | 0.43572 |
| oxidoreduction coenzyme metabolic process (GO:0006733) | 0.00420 | 0.22828 |
| cell development (GO:0048468) | 0.00431 | 0.66789 |
| regulation of carbohydrate catabolic process (GO:0043470) | 0.00438 | 0.22828 |
| negative regulation of cellular component organization (GO:0051129) | 0.00475 | 0.22828 |
| atrial cardiac muscle cell to AV node cell communication (GO:0086066) | 0.00476 | 0.43572 |
| atrial cardiac muscle cell to AV node cell signaling (GO:0086026) | 0.00476 | 0.22828 |
| ion binding (GO:0043167) | 0.00505 | 0.14137 |
| regulation of cofactor metabolic process (GO:0051193) | 0.00643 | 0.25725 |
| nervous system development (GO:0007399) | 0.00661 | 0.43572 |
| gastric acid secretion (GO:0001696) | 0.00720 | 0.43572 |
| negative regulation of reproductive process (GO:2000242) | 0.00807 | 0.43572 |
| excitatory postsynaptic potential (GO:0060079) | 0.00851 | 0.29188 |

Since methylation at different genic features can have different effects on gene expression^45,46^, we also assessed which features were most affected by exposure to maternal diabetes. Relative to all analyzed sites, proportionately fewer loci of interest were found at promoter (8.2% versus 19.5%; hypergeometric test, *p*<1.63×10^-23^) and 5’ UTR regions (4.6% versus 10.8%; *p*<7.25×10^-13^; **Figure 5A**). Conversely, differentially methylated sites were particularly enriched for intronic (28.7% versus 24.5%; *p*<0.001), and intergenic (37.0% versus 25.9%; *p*<3.52×10^-15^) annotations, and less so for coding sequence (16.2% versus 14.4%; *p*<0.05) and non-coding RNA exons (3.4% versus 2.3%; *p*<0.05). These trends were reflected in CpG context annotations, which showed a depletion of sites in CpG islands - commonly found in promoter regions (34.1% versus 37.6%; *p*<0.05), and an enrichment for sites outside of such islands (36.6% versus 34.0%; *p*<0.05; **Figure 5B**). Thus, the majority of significant DNA methylation changes overlapped gene bodies and intergenic regions, which have functional consequences on gene regulation that are more difficult to extrapolate^45,46^.

**Figure 5:**
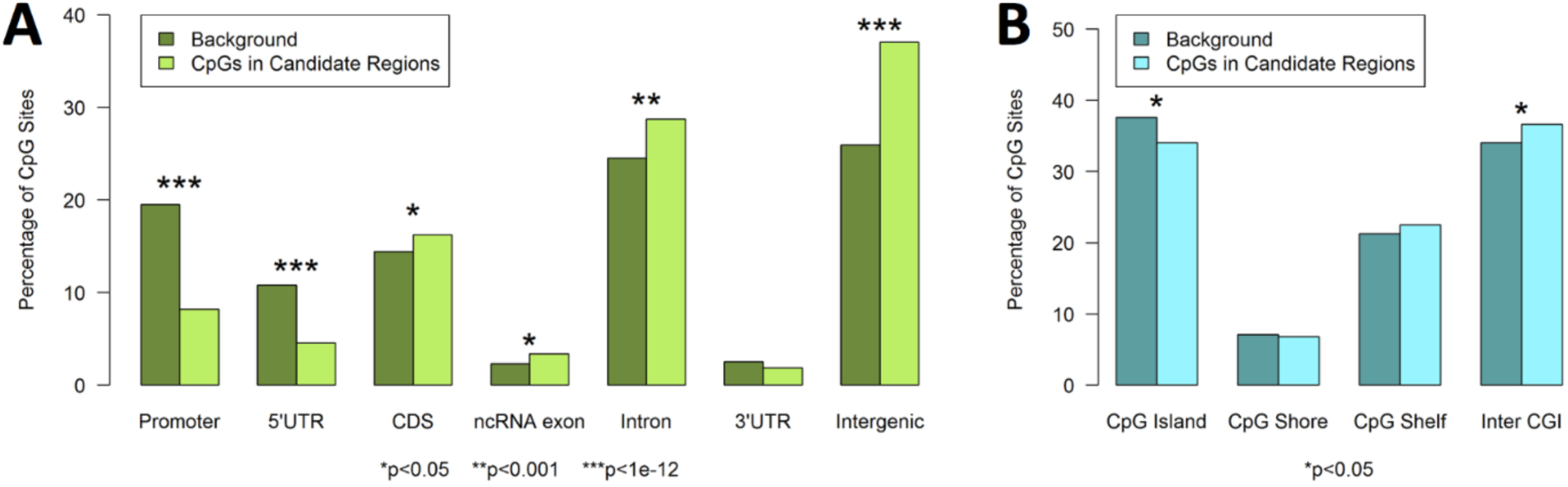
Depletion of differentially methylated sites in promoters accompanied by enrichment of loci in introns and intergenic regions. (**5A**) RefSeq gene annotations. Promoters are defined as the region 2 kb upstream of a transcription start site. UTR=untranslated region; CDS=coding sequence; ncRNA=non-coding RNA. (**5B**) CpG context annotation. CGI=CpG island.

### Allele-specific methylation suggests a role for sequence variation in diabetic embryopathy

Finally, we decided to leverage the genetic information available in our bisulfite sequencing data to evaluate the role of sequence variation in the observed methylation differences. Bisulfite treatment converts unmethylated cytosines to thymines; at the level of single sequencing reads, this can create ambiguity over whether SNVs with C>G, G>C, G>A, or G>T base changes represent true SNVs or artefacts of the bisulfite conversion process. Therefore, we focused on individuals homozygous for both reference and alternate alleles. We identified 541 unique, common SNVs that were homozygous in at least two samples per allele and within 100 bp up- and downstream of 548 unique plus-strand CpG cytosines, resulting in a total of 609 SNV-CpG pairs (**Figure 6A**).

**Figure 6:**
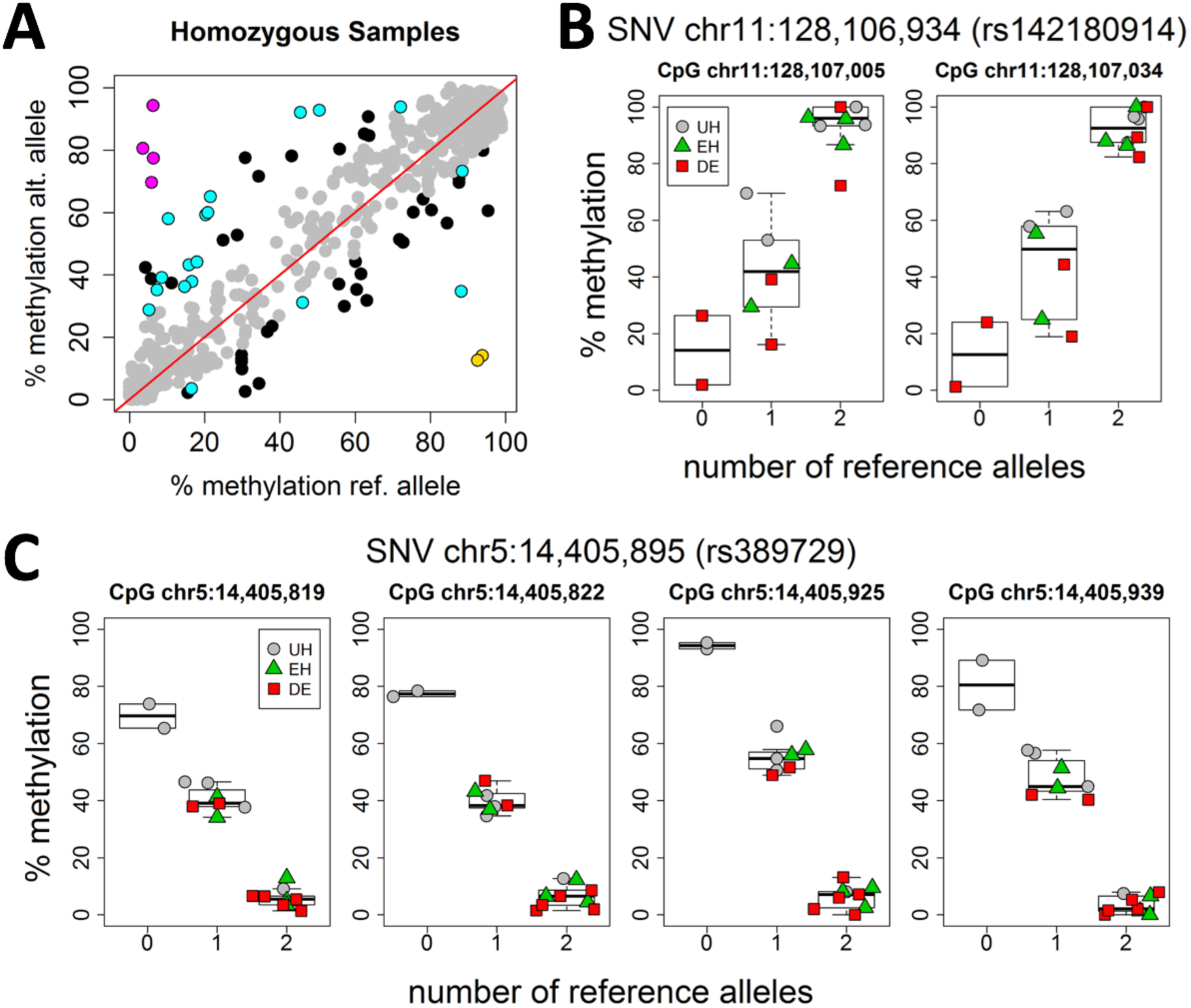
Allele-specific methylation (ASM) at differentially methylated CpG loci. (**6A**) Mean % methylation of homozygous reference samples plotted against homozygous alternate. Each genotype category contains at least two individuals. Colored points are found within clustered regions of differential methylation. Yellow dots show one SNV associated with ASM at two proximal CpG sites found in an intergenic region (**6B**). Pink dots show one SNV associated with ASM at four proximal CpG sites found in *TRIO* (**6C**).

For this exercise, we defined allele-specific methylation (ASM) as a mean difference in the percent methylation associated with the reference vs the alternate SNV allele that fell in the tails (<5^th^ centile; >95^th^ centile) of the distribution. Of the 62 SNV-CpG pairs with evidence of ASM - 24 (39.3%), fell within our candidate bins (**Supplementary Table 6**). These pairs were composed of 13 unique SNVs associated with 23 distinct CpGs and overlapped six RefSeq genes: *CDH12*, *ECHDC3*, *LINC02098*, *NTM*, *RNF157*, and *TRIO.* Among the pairs with the most pronounced difference in percent methylation, were two CpG sites associated with the variant rs142180914 in the lincRNA *LINC02098* that showed increasing DNA methylation with increasing numbers of reference alleles (**Figure 6B**). Remarkably, all samples homozygous for the alternate allele were from infants with DE. We also observed four CpG loci that were strongly correlated with the intronic variant rs389729 in *TRIO*, but showed a decrease in methylation for each additional reference allele (**Figure 6C**). In this case as well, the only individuals homozygous for the alternate allele were diabetes-unexposed controls (**Figure 6C**). Apart from intellectual disability, individuals with heterozygous pathogenic variants in *TRIO* share additional features in common with the phenotypic spectrum of DE, including microcephaly, and digit malformations^47,48^.

## DISCUSSION

The rising prevalence of diabetes and its known teratogenic effects reinforce the need to not only learn more about the disease pathogenesis in IDMs, but to also improve the current diagnosis of diabetic embryopathy. Other studies have already explored the utility of DNA methylation for improved diagnoses of congenital disorders; in Sotos Syndrome (MIM #117550) – a neurodevelopmental disorder associated with tissue overgrowth – loss of function of *NSD1*, a histone methyltransferase, was shown to result in a characteristic genome-wide DNA methylation pattern that could distinguish Sotos patients from controls and individuals with the phenotypically similar Weaver syndrome (MIM #277590)^37^. Likewise, distinct methylation patterns have been proffered in the diagnosis of CHARGE (MIM #214800) and Kabuki (MIM #147920) syndromes^49^. The utility of DNA methylation biomarkers has also been demonstrated in diseases not primarily driven by defects in epigenome maintenance genes, including a range of cancers^50-52^. Similar to our investigation, a recent study of fetal alcohol spectrum disorder – caused by *in utero* exposure to ethanol – also identified significantly altered DNA methylation patterns in buccal epithelial cells^53^.

Our results suggest that, comparable to the aforementioned Mendelian disorders, the teratogenic effect that leads to DE has a large enough influence on offspring DNA methylation to allow accurate and consistent distinction between UH and DE infants using modest sample sizes. More importantly, using PCA, we could also distinguish diabetes-exposed, but healthy infants from diabetes-exposed infants with congenital malformations. Thus, DNA methylation biomarkers may not only be useful for identifying intrauterine diabetes exposure, but may also have the potential to inform the *severity* of teratogenesis. A screening test based on such DNA methylation patterns could be used by clinicians to efficiently evaluate the likelihood of diabetes exposure as the cause of birth defects among IDMs – an option that does not presently exist – and could prove useful for studying the causes of DE in humans. Additionally, a more definitive IDM diagnosis would help to abrogate the emotional and financial costs associated with the prolonged medical odyssey faced by a substantial portion of children with birth defects. This would bring such congenital defects in line with current considerations for the use of whole-exome sequencing in rare Mendelian disorders^54^, thereby furthering appropriate patient-centered care and genetic counseling.

We were careful in our analysis to focus on differential methylation of large magnitude that passed stringent statistical thresholds of significance; however, the sample size employed did not allow us to include covariates such as maternal smoking, which has been associated with altered DNA methylation in offspring^55,56^. Among the DE and UH participants, there was only one instance in each category of reported maternal smoking during pregnancy. The first was an affected (DE) infant who was identified as an outlier and removed from subsequent analyses. The other (UH) infant had an otherwise unremarkable methylation pattern. In our cohort, DE was marginally correlated with maternal obesity, which has been shown to influence offspring DNA methylation^57,58^ and is an independent, albeit less pronounced, risk factor for birth defects^11^. Given that obese women are more likely to develop T2DM^59^ and GDM^60^, it remains to be seen whether the teratogenic mechanisms of maternal obesity are distinct from, or synergistic with, those of maternal diabetes^61^.

For diagnostic purposes, accessible tissue, such as buccal epithelial cells, are sufficient (and even preferred) as long as the characteristic DNA methylation fingerprint of maternal diabetes exposure is consistently correlated with the phenotype. Unlike germline genetic information, however, which is typically the same across all tissues in a given individual, epigenetic information can vary between cell types. Compared to blood – another commonly used and readily-available surrogate tissue – methylation levels in buccal epithelial cells are more consistently correlated with other, non-blood cells^62^, suggesting that our results may closely mirror differential methylation patterns in other organ systems. Formal evaluation of methylation patterns across cell types is needed to fully understand whether the changes in DNA methylation seen in buccal epithelial cells are consistent and, more importantly, causally-implicated in DE.

Elevated glucose concentrations during development, resulting from maternal hyperglycemia, have been shown to amplify glucose metabolism and consequently increase the levels of reactive oxygen species (ROS)^18,63^. DNA damage incurred by ROS is known to prompt cell cycle arrest and induce apoptosis, with the potential to disrupt crucial developmental stages. Nonetheless, a cell that has escaped apoptosis can still be left with substantial damage. Given that double strand break repair can produce heritable changes in DNA methylation at the break locus^64^, it is feasible that the base excision and single strand break repair mechanisms also invoked by ROS damage could result in similar DNA methylation disruptions, which might significantly alter the expression of important developmental genes. Favorably, our gene and pathway analyses converged upon a number of genes affecting embryonic development, including many that underlie known genetic syndromes; this could explain the strong phenotypic overlap between many of these syndromes and DE.

Non-random alterations in DNA methylation patterns could, at least in part, be explained by the influence of genetic factors. By exploring the immediate *cis* DNA sequence of our bisulfite sequencing – a feature not available with conventional methylation arrays – we identified SNVs that correlate with DNA methylation at proximal CpG sites. Differential methylation occurring at CpG sites under the control of *cis*-genetic variation could be the result of differences in the chance distribution of ASM alleles between cases and controls; alternatively, this might reflect genetic variation that, upon exposure to the milieu of maternal diabetes, potentiates changes in DNA methylation in an allele-specific manner and predisposes an individual to DE. Although bisulfite treatment can obscure the state of certain single nucleotide polymorphisms at variable base positions, ever-improving SNV calling algorithms and the adjunctive use of genome sequencing, suggest that this limitation will be surmountable in the near future. As a further advantage of bisulfite sequencing, we were able to evaluate DNA methylation in regions of the genome that are not typically evaluated by microarray-based epigenome screens – 18.6% of our top candidate bins are not represented on the widely-used Illumina450K methylation array. Moreover, sequencing can be done on an individual (rather than batched) level as has been done for rare Mendelian disorders^65,66^, and is thus better suited to the diagnosis of relatively rare teratogenic exposures.

The recent expansion in our understanding of the epigenome has illuminated the formative influence of DNA methylation on human diseases. The assembly of larger IDM cohorts would allow for the diagnostic potential of methylation in IDM to be further refined, enabling an assessment of the impact of previously-mentioned covariates. Recruitment of additional comparison samples, particularly, infants with birth defects but no diabetes exposure, IDMs with less severe complications, and older children/adolescents who are more distant from *in utero* exposure, would allow for robust estimates of the sensitivity and specificity of DNA methylation in the diagnosis of IDM. Finally, the emerging role of DNA methylation across a spectrum of environmental exposures suggests that our approach may be applicable in a variety of other teratogenic exposures, paving the way for long-overdue improvements in diagnostics for this class of disorder.

## ACKNOWLEDGMENTS

This research was supported by a Baylor College of Medicine Presidential Award (JWB) and USDA, ARS cooperative agreement (58-3092-5-001). NAH is funded by a Clinical Scientist Development Award from the Doris Duke Charitable Foundation (Grant #:2013096). KVS is supported by Award Number T32GM008307 from the National Institute of General Medical Sciences. The content is solely the responsibility of the authors and does not necessarily represent the official views of the National Institute of General Medical Sciences or the National Institutes of Health. The authors would like to thank Nancy Hall for invaluable technical help with sample management.

## AUTHOR CONTRIBUTIONS

KVS, KF, JRK, JWB, and NAH designed the study; AB, MSA, and NCS recruited participants and acquired samples; GZ and PH processed and sequenced the samples; KVS, JWB, and NAH directed data analyses; KVS and NAH wrote the paper; All authors read and edited the manuscript.

## REFERENCES

1. Wilson, J. G., Environment and Birth Defects (Academic Press, New York, 1973).

2. Brent, R. L., The cause and prevention of human birth defects: What have we learned in the past 50 years? Congenit.l Anom. (Kyoto) 41 (1), 3-21 (2001).

3. Mattson, S. N., Crocker, N. & Nguyen, T. T., Fetal Alcohol Spectrum Disorders: Neuropsychological and Behavioral Features. Neuropsychol. Rev. 21 (2), 81-101 (2011).

4. Vajda, F. J. et al., Critical relationship between sodium valproate dose and human teratogenicity: results of the Australian register of anti-epileptic drugs in pregnancy. J. Clin. Neurosci. 11 (8), 854-858 (2004).

5. Harden, C. L. et al., Practice Parameter update: Management issues for women with epilepsy-Focus on pregnancy (an evidence-based review): Teratogenesis and perinatal outcomes Report of the Quality Standards Subcommittee and Therapeutics and Technology Assessment Subcommittee of the American Academy of Neurology and American Epilepsy Society. Neurology 73 (2), 133-141 (2009).

6. Sheffield, J. S., Butler-Koster, E. L., Casey, B. M., McIntire, D. D. & Leveno, K. J., Maternal Diabetes Mellitus and Infant Malformations. Obstet. Gynecol. 100 (5, Part 1), 925-930 (2002).

7. Lepercq, J., French Multicentric Survey of Outcome of Pregnancy in Women With Pregestational Diabetes. Diabetes Care 26 (11), 2990-2993 (2003).

8. Evers, I. M., de Valk, H. W. & Visser, G. H. A., Risk of complications of pregnancy in women with type 1 diabetes: nationwide prospective study in the Netherlands. BMJ 328, 915 (2004).

9. Sharpe, P. B., Chan, A., Haan, E. A. & Hiller, J. E., Maternal diabetes and congenital anomalies in South Australia 1986–2000: A population-based cohort study. Birth Defects Res. A Clin. Mol. Teratol. 73 (9), 605-611 (2005).

10. Jonasson, J. M. et al., Fertility in Women With Type 1 Diabetes: A population-based cohort study in Sweden. Diabetes Care 30 (9), 2271-2276 (2007).

11. Block, S. R. et al., Maternal Pre-Pregnancy Body Mass Index and Risk of Selected Birth Defects: Evidence of a Dose-Response Relationship. Paediatr. Perinat. Epidemiol. 27 (6), 521-531 (2013).

12. Persson, M. et al., Risk of major congenital malformations in relation to maternal overweight and obesity severity: cohort study of 1.2 million singletons. BMJ 357, j2563 (2017).

13. García-Patterson, A. et al., In human gestational diabetes mellitus congenital malformations are related to pre-pregnancy body mass index and to severity of diabetes. Diabetologia 47 (3), 509-514 (2004).

14. Schaefer-Graf, U. M. et al., Patterns of congenital anomalies and relationship to initial maternal fasting glucose levels in pregnancies complicated by type 2 and gestational diabetes. Am. J. Obstet. Gynecol. 182 (2), 313-320 (2000).

15. Shields, L. E., Gan, E. A., Murphy, H. F., Sahn, D. J. & Moore, T. R., The Prognostic Value of Hemoglobin A1c in Predicting Fetal Heart Disease in Diabetic Pregnancies. Obstet. Gynecol. 81 (6), 954-957 (1993).

16. Ylinen, K., Aula, P., Stenman, U.-H., Kesäniemi-Kuokkanen, T. & Teramo, K., Risk of minor and major fetal malformations in diabetics with high haemoglobin A1c values in early pregnancy. Br Med J (Clin Res Ed) 289 (6441), 345-346 (1984).

17. Dominguez-Salas, P. et al., Maternal nutrition at conception modulates DNA methylation of human metastable epialleles. Nat. Commun. 5, 3746 (2014).

18. Wei, D. & Loeken, M. R., Increased DNA Methyltransferase 3b (Dnmt3b)-Mediated CpG Island Methylation Stimulated by Oxidative Stress Inhibits Expression of a Gene Required for Neural Tube and Neural Crest Development in Diabetic Pregnancy. Diabetes 63 (10), 3512-3522 (2014).

19. Bouchard, L. et al., Leptin Gene Epigenetic Adaption to Impaired Glucose Metabolism During Pregnancy. Diabetes Care 33 (11), 2436-2441 (2010).

20. Bouchard, L. et al., Placental Adiponectin Gene DNA Methylation Levels Are Associated With Mothers’ Blood Glucose Concentration. Diabetes 61 (5), 1272-1280 (2012).

21. El Hajj, N. et al., Metabolic Programming of *MEST* DNA Methylation by Intrauterine Exposure to Gestational Diabetes Mellitus. Diabetes 62 (4), 1320-1328 (2013).

22. Allard, C. et al., Mendelian randomization supports causality between maternal hyperglycemia and epigenetic regulation of leptin gene in newborns. Epigenetics 10 (4), 342-351 (2015).

23. Ruchat, S.-M. et al., Gestational diabetes mellitus epigenetically affects genes predominantly involved in metabolic diseases. Epigenetics 8 (9), 935-943 (2013).

24. Petropoulos, S. et al., Gestational Diabetes Alters Offspring DNA Methylation Profiles in Human and Rat: Identification of Key Pathways Involved in Endocrine System Disorders, Insulin Signaling, Diabetes Signaling, and ILK Signaling. Endocrinology 156 (6), 2222-2238 (2014).

25. Finer, S. et al., Maternal gestational diabetes is associated with genome-wide DNA methylation variation in placenta and cord blood of exposed offspring. Hum. Mol. Genet. 24 (11), 3021-3029 (2015).

26. Haertle, L. et al., Epigenetic signatures of gestational diabetes mellitus on cord blood methylation. Clinical Epigenetics 9, 28 (2017).

27. West, N. A., Kechris, K. & Dabelea, D., Exposure to Maternal Diabetes in Utero and DNA Methylation Patterns in the Offspring. Immunometabolism 1, 1-9 (2013).

28. Gautier, J.-F. et al., Kidney Dysfunction in Adult Offspring Exposed In Utero to Type 1 Diabetes Is Associated with Alterations in Genome-Wide DNA Methylation. PLoS One 10 (8), e0134654 (2015).

29. del Rosario, M. C. et al., Potential epigenetic dysregulation of genes associated with MODY and type 2 diabetes in humans exposed to a diabetic intrauterine environment: An analysis of genome-wide DNA methylation. Metabolism 63 (5), 654-660 (2014).

30. Reinius, L. E. et al., Differential DNA Methylation in Purified Human Blood Cells: Implications for Cell Lineage and Studies on Disease Susceptibility. PLoS One 7 (7), e41361 (2012).

31. Krueger, F. & Andrews, S. R., Bismark: a flexible aligner and methylation caller for Bisulfite-Seq applications. Bioinformatics 27 (11), 1571-1572 (2011).

32. Hicks, S. C. & Irizarry, R. A., quantro: a data-driven approach to guide the choice of an appropriate normalization method. Genome Biol. 16, 117 (2015).

33. Horvath, S., DNA methylation age of human tissues and cell types. Genome Biol. 14, R115 (2013).

34. Akalin, A. et al., methylKit: a comprehensive R package for the analysis of genome-wide DNA methylation profiles. Genome Biol. 13, R87 (2012).

35. Park, Y., Figueroa, M. E., Rozek, L. S. & Sartor, M. A., MethylSig: a whole genome DNA methylation analysis pipeline. Bioinformatics, 30 (17), 2414-2422 (2014).

36. Hall, M. et al., The WEKA data mining software: an update. ACM SIGKDD Explorations Newsletter 11 (1), 10-18 (2009).

37. Choufani, S. et al., NSD1 mutations generate a genome-wide DNA methylation signature. Nat. Commun. 6, 10207 (2015).

38. Kamburov, A., Stelzl, U., Lehrach, H. & Herwig, R., The ConsensusPathDB interaction database: 2013 update. Nucleic Acids Res. 41 (D1), D793-800 (2013).

39. Liu, Y., Siegmund, K. D., Laird, P. W. & Berman, B. P., Bis-SNP: Combined DNA methylation and SNP calling for Bisulfite-seq data. Genome Biol. 13, R61 (2012).

40. McKenna, A. et al., The Genome Analysis Toolkit: A MapReduce framework for analyzing next-generation DNA sequencing data. Genome Res. 20 (9), 1297-1303 (2010).

41. Elliott, G. et al., Intermediate DNA methylation is a conserved signature of genome regulation. Nat. Commun. 6, 6363 (2015).

42. Harada, M. et al., G-CSF prevents cardiac remodeling after myocardial infarction by activating the Jak-Stat pathway in cardiomyocytes. Nat. Med. 11 (3), 305-311 (2005).

43. Miletich, I. et al., Developmental stalling and organ-autonomous regulation of morphogenesis. Proc. Natl. Acad. Sci. USA 108 (48), 19270-19275 (2011).

44. Iwashita, S. et al., Versatile Roles of R-Ras GAP in Neurite Formation of PC12 Cells and Embryonic Vascular Development. J. Biol. Chem. 282 (6), 3413-3417 (2007).

45. Neri, F. et al., Intragenic DNA methylation prevents spurious transcription initiation. Nature 543 (7643), 72-77 (2017).

46. Wong, J. J.-L. et al., Intron retention is regulated by altered MeCP2-mediated splicing factor recruitment. Nat. Commun. 8, 15134 (2017).

47. Ba, W. et al., TRIO loss of function is associated with mild intellectual disability and affects dendritic branching and synapse function. Hum. Mol. Genet. 25 (5), 892-902 (2015).

48. Pengelly, R. J. et al., Mutations specific to the Rac-GEF domain of *TRIO* cause intellectual disability and microcephaly. J. Med. Genet. 53 (11), 735-742 (2016).

49. Butcher, D. T. et al., CHARGE and Kabuki Syndromes: Gene-Specific DNA Methylation Signatures Identify Epigenetic Mechanisms Linking These Clinically Overlapping Conditions. Am. J. Hum. Genet. 100 (5), 773-788 (2017).

50. Chung, W. et al., Detection of Bladder Cancer Using Novel DNA Methylation Biomarkers in Urine Sediments. Cancer Epidemiol. Biomarkers Prev. 20 (7), 1483-1491 (2011).

51. de Martino, M., Klatte, T., Haitel, A. & Marberger, M., Serum Cell-Free DNA in Renal Cell Carcinoma: A Diagnostic and Prognostic Marker. Cancer 118 (1), 82-90 (2012).

52. Payne, S. E., From discovery to the clinic: the novel DNA methylation biomarker (m)SEPT9 for the detection of colorectal cancer in blood. Epigenomics 2 (4), 575-585 (2010).

53. Portales-Casamar, E. et al., DNA methylation signature of human fetal alcohol spectrum disorder. Epigenet. Chromatin 9, 25 (2016).

54. Stark, Z. et al., Prospective comparison of the cost-effectiveness of clinical whole-exome sequencing with that of usual care overwhelmingly supports early use and reimbursement. Genet. Med., doi:10.1038/gim.2016.221 (2017).

55. Breton, C. V. et al., Prenatal Tobacco Smoke Exposure Affects Global and Gene-specific DNA Methylation. Am. J. Respir. Crit. Care Med. 180 (5), 462-467 (2009).

56. Richmond, R. C. et al., Prenatal exposure to maternal smoking and offspring DNA methylation across the lifecourse: findings from the Avon Longitudinal Study of Parents and Children (ALSPAC). Hum. Mol. Genet. 24 (8), 2201-2217 (2014).

57. Liu, X. et al., Maternal Preconception Body Mass Index and Offspring Cord Blood DNA Methylation: Exploration of Early Life Origins of Disease. Environ. Mol. Mutagen. 55 (3), 223-230 (2014).

58. Sharp, G. C. et al., Maternal pre-pregnancy BMI and gestational weight gain, offspring DNA methylation and later offspring adiposity: findings from the Avon Longitudinal Study of Parents and Children. Int. J. Epidemiol. 44 (4), 1288-1304 (2015).

59. Colditz, G. A. et al., Weight as a risk factor for clinical diabetes in women. Am. J. Epidemiol. 132 (3), 501-513 (1990).

60. Chu, S. Y. et al., Maternal Obesity and Risk of Gestational Diabetes Mellitus. Diabetes Care 30 (8), 2070-2076 (2007).

61. Anderson, J. L. et al., Maternal Obesity, Gestational Diabetes, and Central Nervous System Birth Defects. Epidemiology 16 (1), 87-92 (2005).

62. Lowe, R. et al., Buccals are likely to be a more informative surrogate tissue than blood for epigenome-wide association studies. Epigenetics 8 (4), 445-454 (2013).

63. Zabihi, S. & Loeken, M. R., Understanding Diabetic Teratogenesis: Where Are We Now and Where Are We Going? Birth Defects Res. A Clin. Mol. Teratol. 88 (10), 779-790 (2010).

64. Russo, G. et al., DNA damage and Repair Modify DNA methylation and Chromatin Domain of the Targeted Locus: Mechanism of allele methylation polymorphism. Sci. Rep. 6, 33222 (2016).

65. Yang, Y. et al., Molecular Findings Among Patients Referred for Clinical Whole-Exome Sequencing. JAMA 312 (18), 1870-1879 (2014).

66. Yang, Y. et al., Clinical Whole-Exome Sequencing for the Diagnosis of Mendelian Disorders. N. Engl. J. Med. 369 (16), 1502-1511 (2013).

